# Sensory regulation of absence seizures in a mouse model of *Gnb1* encephalopathy

**DOI:** 10.1101/2022.04.28.489780

**Authors:** Sasa Teng, Fenghua Zhen, Briana R. McRae, Elaine Zhu, Wayne N. Frankel, Yueqing Peng

## Abstract

Absence seizures, a type of non-convulsive epilepsy manifested by spike-wave discharges (SWD) in the electroencephalogram (EEG), display synchronous reciprocal excitation between the neocortex and thalamus. Recent studies have revealed that inhibitory neurons in the reticular thalamic (RT) nucleus and excitatory thalamocortical (TC) neurons are two key subcortical players in generating SWD. However, the signals that drive SWD-related thalamic activity remain elusive. Here, we show that SWD predominately occurs during wakefulness in several mouse models of absence epilepsy. In more focused studies of *Gnb1* mutant mice, we found that sensory input regulates SWD. Using *in vivo* recording, we demonstrate that TC cells are activated prior to the onset of SWD and then inhibited during SWD. On the contrary, RT cells are slightly inhibited prior to SWD, but are strongly activated during SWD. Furthermore, chemogenetic activation of TC cells leads to the enhancement of SWD in Gnb1 mice. Together, our results indicate that sensory input in the periphery can regulate SWD by activating the thalamocortical pathway.

## INTRODUCTION

Spike-wave discharges (SWD) are the electroencephalographic (EEG) hallmarks of absence seizures, a type of generalized seizures commonly observed in many neurodevelopmental disorders. Growing evidence has supported that the thalamocortical circuits act as the primary generator of the SWD^1,2^. The concept originated from a hypothesis that SWD may develop by the same thalamocortical circuits which normally create sleep spindles under certain conditions of cortical hyperexcitability^3,4^. In particular, GABAergic neurons in the reticular thalamic (RT) nucleus, excitatory thalamocortical (TC) cells, and neocortical pyramidal cells comprise a circuit that sustains the thalamocortical oscillatory firing of absence seizures^5–7^. Despite the well-characterized circuit mechanisms underlying the SWD, little is known how the seizure events start, or what signals trigger the seizures. One of the limitations is that most circuit studies were performed *in vitro*^8,9^ or *in silico*^10^, which lack behavioral relevance to SWD. Here, we use *in vivo* recording and neural manipulation to understand the neural signals that drive the absence seizures in a mouse model of *Gnb1* encephalopathy.

Recent clinic studies show that mutations in *Gnb1*, encoding the Gβ1 subunit of G proteins, cause GNB1 encephalopathy, a severe neurodevelopmental disorder characterized by global developmental delay, speech and ambulatory deficits, intellectual disability, and a variety of seizure types^11,12^. Using CRISPR/Cas9, a mouse model of the K78R human pathogenic variant was generated to elucidate how GNB1 mutations cause disease^13^. The Gnb1^K78R/+^ mice recapitulate many clinical features of affected individuals, including developmental delay, motor and cognitive deficits, and absence-like generalized seizures^13^. Here, we use the Gnb1^K78R/+^ mouse model to examine the thalamocortical circuits underlying absence seizures.

In this study, we firstly performed EEG-video recordings in *Gnb1* mutant mice and demonstrated that the brain states strongly regulate the occurrence of SWD. This result has been confirmed in other mouse models of absence epilepsy. Next, we found that sensory input during wake periods can increase SWD in *Gnb1* mice. Using fiber photometry-EEG recordings, we then observed that RT cells are activated, and TC cells are inhibited during SWD. Finally, using chemogenetic manipulation, we further demonstrated that activation of TC cells is sufficient to enhance the SWD in *Gnb1* mice. Taken together, these results indicate that by activating the thalamocortical circuits, sensory activity during wakefulness might drive the absence seizures under pathological condition.

## RESULTS

### Brain states regulates SWD

To identify neural signals that drive the seizure events, we first examined the timing of SWD occurrence. A prior study in GAERS rats shows that vigilance states can predict SWD on a short time scale^14^. To study the relationship between brain states and SWD, we performed EEG and EMG recordings in Gnb1^K78R/+^ mutant and control mice and analyzed their sleep patterns and seizure events. Consistent with our prior study^13^, we observed very frequent SWD events (1597 ± 185 events per 24 hours) with a peak frequency of ~7 Hz in Gnb1^K78R/+^ mutant mice (**Supplementary Fig. 1A-D**). By correlating SWD and wake/sleep states, we found that SWD mostly occur during wakefulness (**Figure 1A-B**). To quantify their correlation, we aligned all SWD events to their onset and examined brain states prior to the seizures. We found that the majority of SWD events occur during wakefulness (83.46% ± 0.86%, mean ± SEM) but not during non-rapid-eye-movement (NREM) sleep (2.07% ± 0.68%) or REM sleep (2.68% ± 0.38%, **Figure 1C, 1E**). Interestingly, we observed a small increase of NREM sleep following the SWD (**Figure 1C**), suggesting that SWD likely affect subsequent brain states. While aligning SWD and EMG data, we noticed that EMG amplitude was decreased during the SWD, compared to that before and after SWD (**Figure 1D, 1F**). The EMG drop suggests reduced locomotor activity during SWD. As reported, frequent SWD in rodent models are indicative of absence seizures^15–17^, which are usually accompanied by behavioral arrest^18–20^. To quantify animal behavior in *Gnb1* mutant mice, we used an infrared camera synchronized to the EEG with automated tracking to record animal movement during absence seizure (**Supplementary Fig. 1E**). We observed decreased locomotion during SWD, particularly during the periods with frequent SWD (**Supplementary Fig. 1F**). To quantify the relationship between animal movement and the intensity of SWD, we calculated locomotion during SWD and grouped them with different durations or different inter-SWD intervals. We found that animal’s movement was inversely correlated with SWD durations, and positively correlated with inter-SWD intervals (**Supplementary Fig. 1G**). These results demonstrated that longer or more frequent SWD are more likely associated with behavioral arrest during absence seizures.

**Figure 1.**
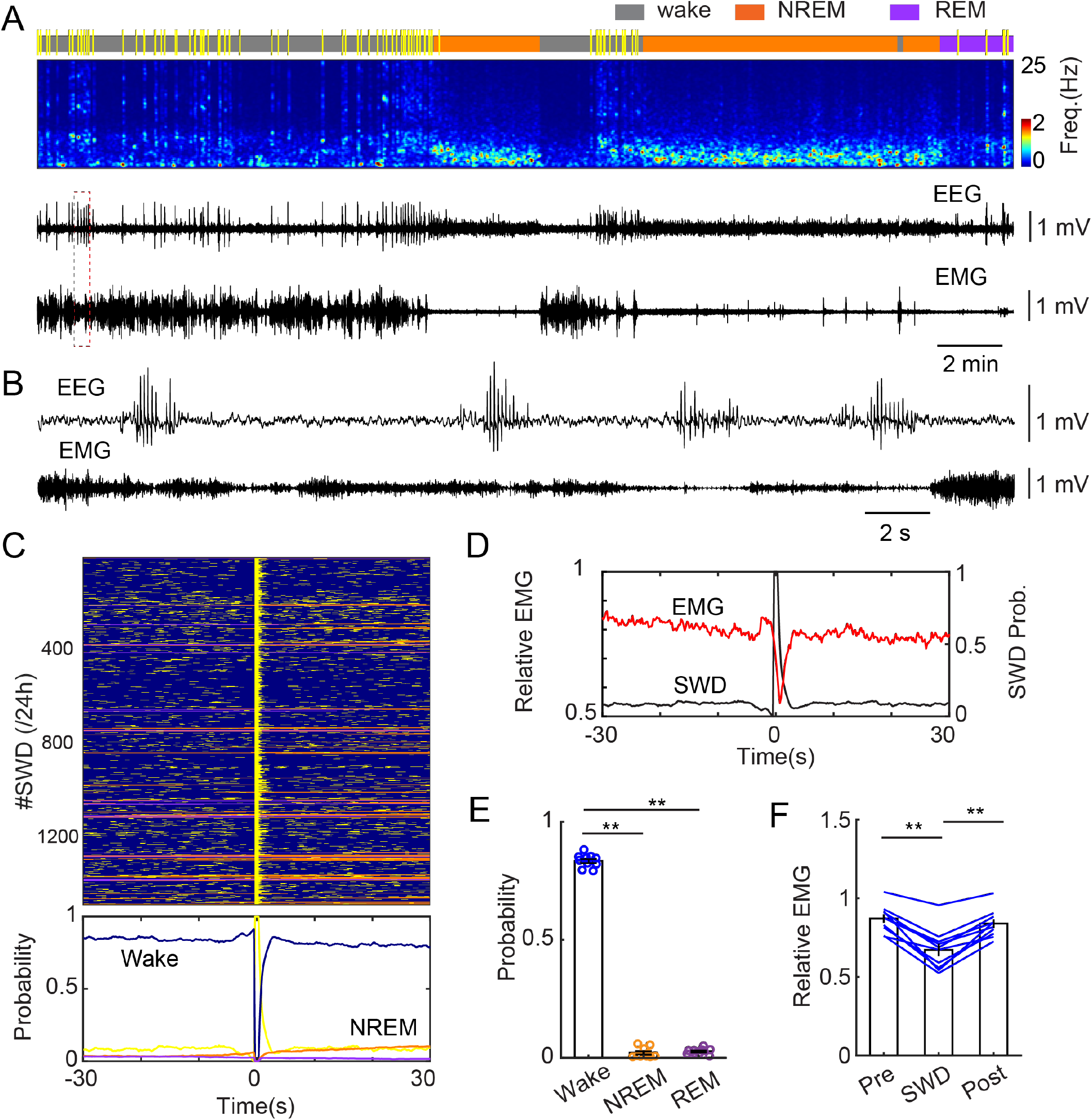
Brain states regulate absence seizures in Gnb1 mice. **A**. A representative recording session in a Gnb1 heterozygous mouse. From top to bottom: brain states (yellow for SWD), EEG spectrogram (0-25 Hz), EEG trace, and EMG trace. EEG and EMG in the red dashed-line box enlarged in **B**. **C**, Top, Brain states (blue for wake, orange for NREM, purple for REM, yellow for SWD) aligned to the onset of 1479 SWD events captured over 24 hours in a Gnb1 mouse. Bottom, probability of brain states before, during, and after SWD events. **D**, Relative EMG amplitude (red) before, during, and after the same SWD events shown in C. The probability of SWD events shown in black. Time 0 indicates the onset of SWD events. **E**, Quantitation of probability of wake, NREM, and REM sleep before the SWD onsets (N = 10 Gnb1 mice, ** P<0.01, paired t-test). **F**. Quantitation of relative EMG amplitude before (pre), during (SWD), and after (post) SWD events (N = 10 Gnb1 mice, ** P<0.01, paired t-test).

The wake-prevalence of SWD is not due to the mice spending more time in wake than in sleep states. Indeed, over a 24 h period, WT and Gnb1^K78R/+^ mice spend a similar amount of total time in wake, NREM and REM sleep (**Supplementary Fig. 2**). Unexpectedly, the Gnb1 mutant mice show flattened circadian patterns of wake/sleep cycles. In particular, the mutant mice show increased wake time, decreased NREM and REM sleep during the light phase, and reversed patterns during the dark phase (**Supplementary Fig. 2**). These results indicated disrupted circadian rhythm in Gnb1 mutant mice.

To test if the correlation between SWD and wakefulness is a general phenomenon in mouse absence seizures, we performed same EEG recording and analysis in other two mouse models that display spontaneous SWD. Firstly, we performed EEG recording in *Stxbp1*-haplodeficient (*Stxbp1^+/-^*) mice, which represent mutations in STXBP1 (Syntaxin-binding protein 1, also known as MUNC18-1), a presynaptic protein essential for neurotransmitter release^21,22^. Heterozygous mutations in STXBP1 have been linked to various severe early epileptic encephalopathies and neurodevelopmental disorders^23,24^. Correlation analysis between SWD and sleep in *Stxbp1*^+/-^ mice demonstrated that SWD predominately occur during wakefulness (78.26% ± 4.16%) but not during sleep states (NREM sleep: 5.25% ± 2.21%; REM sleep: 12.08% ± 4.62%; **Figure 2A-C**). Then, we repeated experiments in *Gria4* deficient (*Gria4^-/-^*) mice, which display spontaneous SWD^25,26^. *GRIA4* encodes a glutamate AMPA receptor subunit known as GluR4. A similar correlation between SWD and wakefulness (84.51 ± 1.75%) was observed in *Gria4*^-/-^ mice (**Figure 2D-F**). In addition, we also observed increased probability of NREM sleep following the SWD in both mouse models (**Figure 2A, 2D**). Taken together, our results indicate that brain states strongly regulate the occurrence of SWD in absence epilepsies.

**Figure 2.**
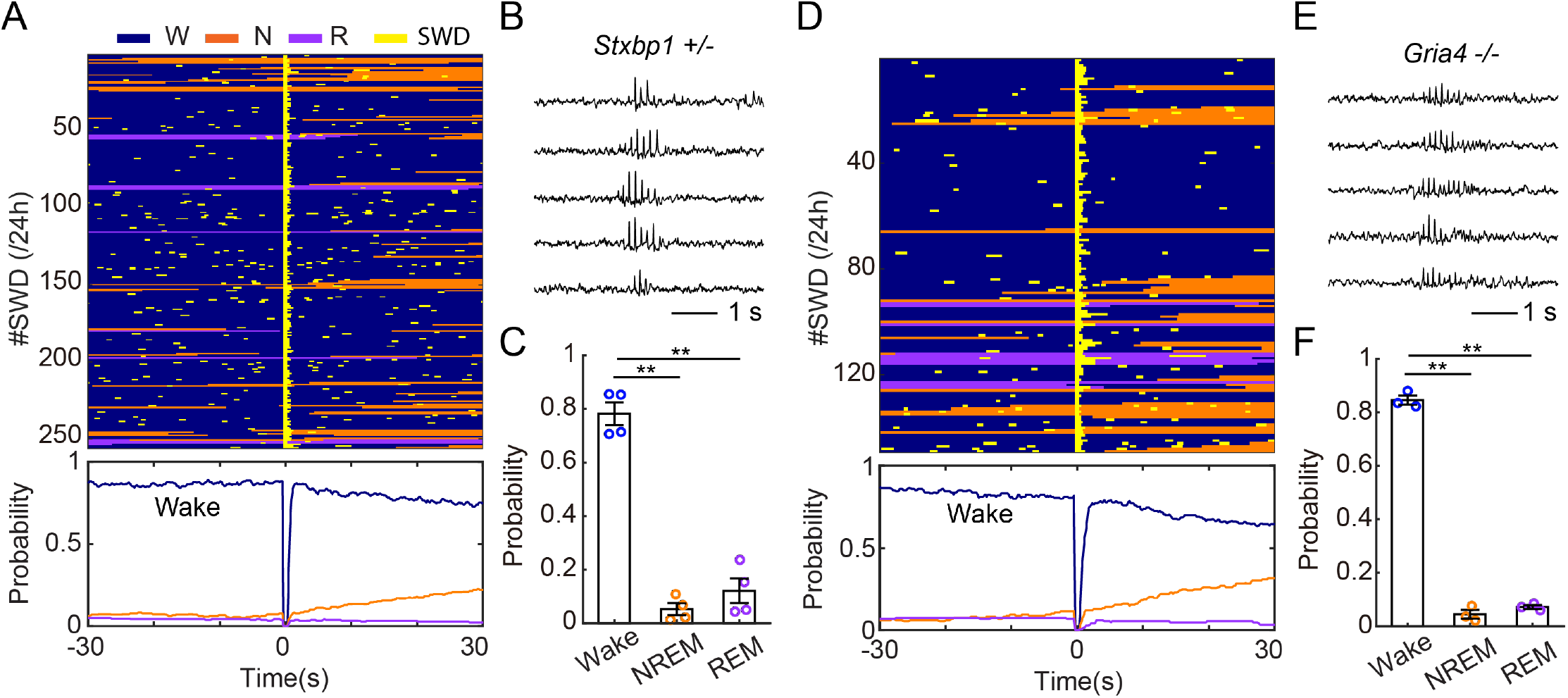
Brain states regulate SWD in other mouse models of absence epilepsy. **A**, Top, Brain states aligned to the onset of SWD events captured over 24 hours in a Stxbp1^+/-^ mouse. Bottom, probability of brain states before, during, and after SWD events. **B**, Representative EEG traces of SWD in the Stxbp1^+/-^ mouse. **C**, Quantitation of probability of wake, NREM, and REM sleep before the SWD onsets in Stxbp1^+/-^ mice (N = 4, ** P<0.01, paired t-test). **D**, Top, Brain states aligned to the onset of SWD events captured over 24 hours in a Gria4^-/-^ mouse. Bottom, probability of brain states before, during, and after SWD events. **E**, Representative EEG traces of SWD in the Gria4^-/-^ mouse. **F**, Quantitation of probability of wake, NREM, and REM sleep before the SWD onsets in Gria4^-/-^ mice (N = 3, ** P<0.01, paired t-test).

### Sensory input regulates SWD in Gnb1 mice

Next, we sought to identify the neural signals that drive or facilitate the SWD in mutant mice. Given the fact that SWD predominately occur during wakefulness, we hypothesized that sensory input from the periphery to the brain during wake periods can regulate SWD. To test this idea, we applied gentle air puffs to the Gnb1 mutant animals and examine the effect on the SWD (**Figure 3A-B**). To quantify, we calculated the number and duration of SWD before, during, and after stimulation in each session. As shown in **Figure 3C**, sensory input significantly increased the number and the durations of SWD during the stimulation period, compared to the period before and after treatment. To further study the sensory effect on SWD over a short time scale, we aligned SWD events to the onset of stimulation in each trial. We observed a decrease of SWD incidence upon the onset of stimulation, followed by a gradual increase during the stimulation period (**Figure 3D**). The dynamic process of SWD changes implies complicated neural activity in the brain. Together, our results indicate that sensory input can regulate SWD in Gnb1 mice.

**Figure 3.**
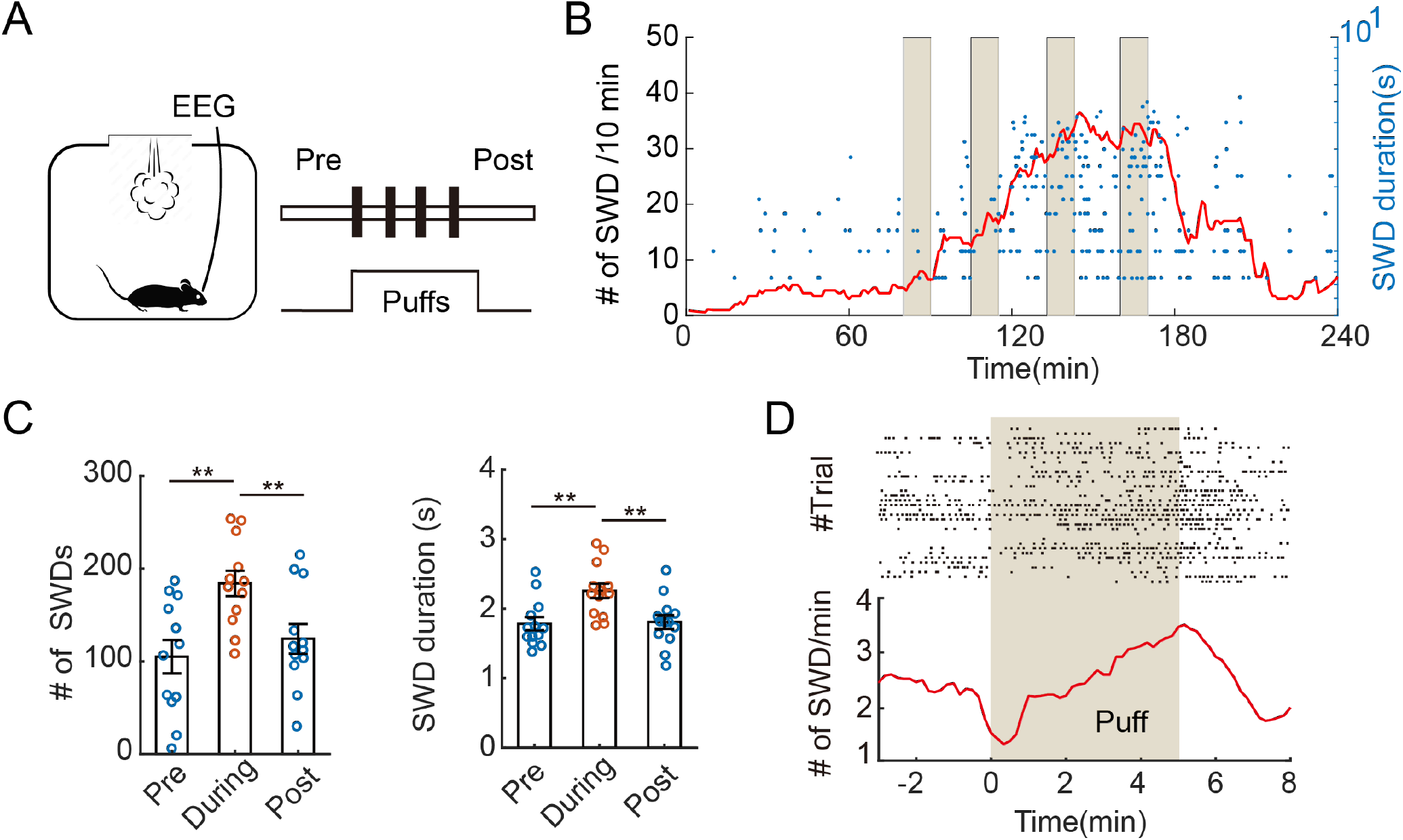
Sensory input regulates absence seizures in Gnb1 mice. **A,** Schematic of experimental design. **B,** A representative session showing the effect of sensory stimulation (four trials of air puffs, shade) on the occurrence of SWD events in a Gnb1 mutant mouse. **C**, Quantitation of SWD number (Left) and duration (Right) before (pre), during, and after (post) sensory stimulation. SWD number was normalized to number per hour during wakefulness. Data were acquired from 13 behavioral sessions in 10 Gnb1 mutant mice (** P<0.01, paired t-test). **D**, Top, raster plot of SWD events aligned to the onset of air puffs. Bottom, peristimulus time histograms (PSTH) of SWD number per minute before, during, and after stimulation. The shade indicates the period of sensory stimulation. Data obtained from 33 trials in 10 Gnb1 mutant mice.

### Thalamic activity is correlated with SWD in Gnb1 mice

Thalamocortical circuits have been implied in the absence seizure in humans^27^ and the SWD in several mouse models^2,28^, including *Scn8a*^29^ and *Stxbp1*^30^. We reasoned that sensory inputs facilitate SWD in mutant mice by activating thalamocortical (TC) cells. To examine thalamic activity associated with SWD, we performed *in vivo* recording in Gnb1 mutant mice by using fiber photometry. Specifically, we stereotaxically injected AAV-CaMKII-GCaMP6s and implant an optic fiber in the sensory part of the thalamus (ventral posteromedial nucleus, or VPM) in Gnb1^+/-^ mice (**Figure 4A**). EEG and EMG electrodes were also implanted during surgery to detect seizures and wake/sleep states. Two-weeks after recovery, we concurrently recorded EEG, EMG, and photometric signals in freely moving animals. As expected, we observed increased spontaneous activity during wakefulness, compared to that in NREM sleep, indicative of sensory activity during wake periods (**Figure 4B**, **Supplementary Fig. 3A-B**). The thalamic activity during REM sleep is largely similar to that during wakefulness, but with variations among different episodes (**Supplementary Fig. 3B**). Strikingly, we found that neural activity in TC cells was significantly suppressed during SWD (**Figure 4C-E**). This inhibitory response was repeatable across different trials (**Figure 4D**) and different animals (**Figure 4E**). Notably, we also observed increased activity of TC cells in the 2-3s time window prior to the SWD (**Figure 4E**), which might suggest the role of thalamic activity in the induction of SWD.

**Figure 4.**
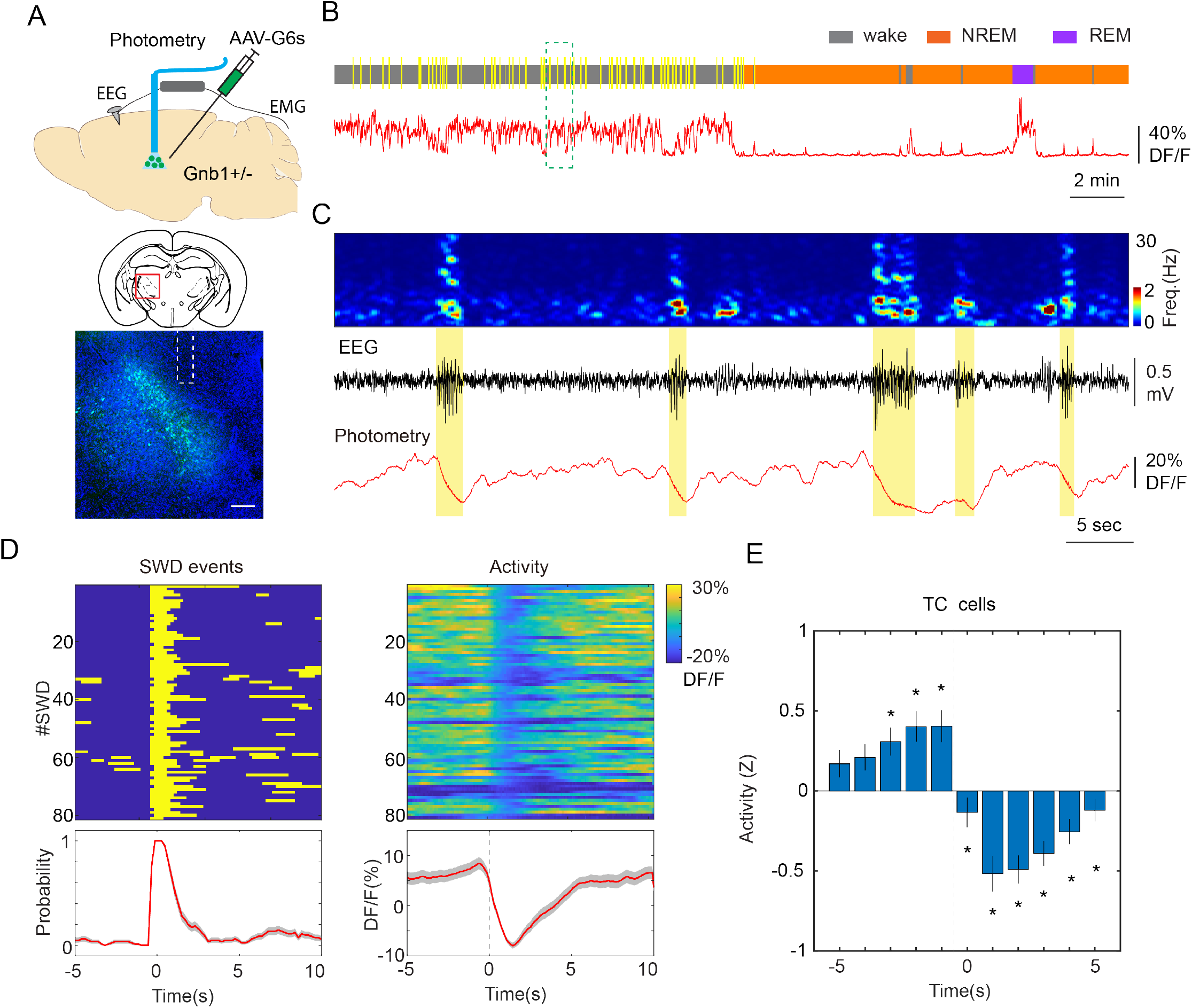
Activity of thalamic cells during SWD events in Gnb1 mice. **A,** Schematic of experimental design. AAV-CaMKII-GCaMP6s was unilaterally injected in the thalamus of Gnb1^+/-^ mice. An optic fiber was implanted above the injection site. Middle, atlas of a brain section (AP −1.82). Bottom, fluorescence image showing GCaMP6s expression (green) in the thalamus (red box above). The white dash line indicates the placement of optic fiber. Blue, DAPI. Scale bar, 200 um. **B,** A representative recording session showing neural activity in thalamic cells of a Gnb1 mutant mouse during wake and sleep states (gray, wake; orange, NREM sleep; purple, REM sleep; yellow, SWD). **C**, Enlarged view (green dash-line box above) showing TC activity during SWD. From top to bottom, EEG spectrogram (0-30 Hz), EEG trace, and photometric signal. **D**, Left, aligned SWD events in a recording session (1 hour). Right top, color-coded calcium signals aligned to the onset of SWD events in the recording session shown on Left. Right bottom, averaged calcium signals across trials. Shading, SEM. **E**, Quantitation of calcium activity of TC cells before, during, and after SWD events. Datapoints at Time −5 were used as baseline for statistical comparison. The dash line indicates the onset of SWD events. Data represent Z values, normalized to the average DF/F during wake periods (12 recording sessions from 4 Gnb1^+/-^ animals, * P<0.001, paired t-test).

Next, we examined the activity in the reticular thalamic nucleus (RT). Previous studies show that the RT is critical for the generation of SWD. For instance, Makinson et al. showed that selective knockdown of *Scn8a* in RT is sufficient to generate the SWD^29^. To genetically target RT cells, we crossed Gnb1 mutant mice with PV-Cre transgenic mice (as reported^29^) to generate Gnb1^+/-^;PV-Cre mice. Then, we stereotaxically injected AAV-FLEX-GCaMP6s in the RT and implanted an optic fiber above the injection site (**Figure 5A**). Two weeks after recovery, we performed photometry-EEG recordings in Gnb1^+/-^;PV-Cre mice. Opposite to TC activity, we observed increased activity in RT cells during NREM sleep, compared to wakefulness (**Figure 5B, Supplementary Fig. 3C-D**). The RT activity during REM sleep was inhibited (**Supplementary Fig. 3D**). More importantly, we found that the SWD reliably evoked calcium increase in RT cells (**Figure 5C-E**), indicating that RT cells are active during SWD. Interestingly, the activity of RT cells was slightly decreased prior to the SWD (**Figure 5E**).

**Figure 5.**
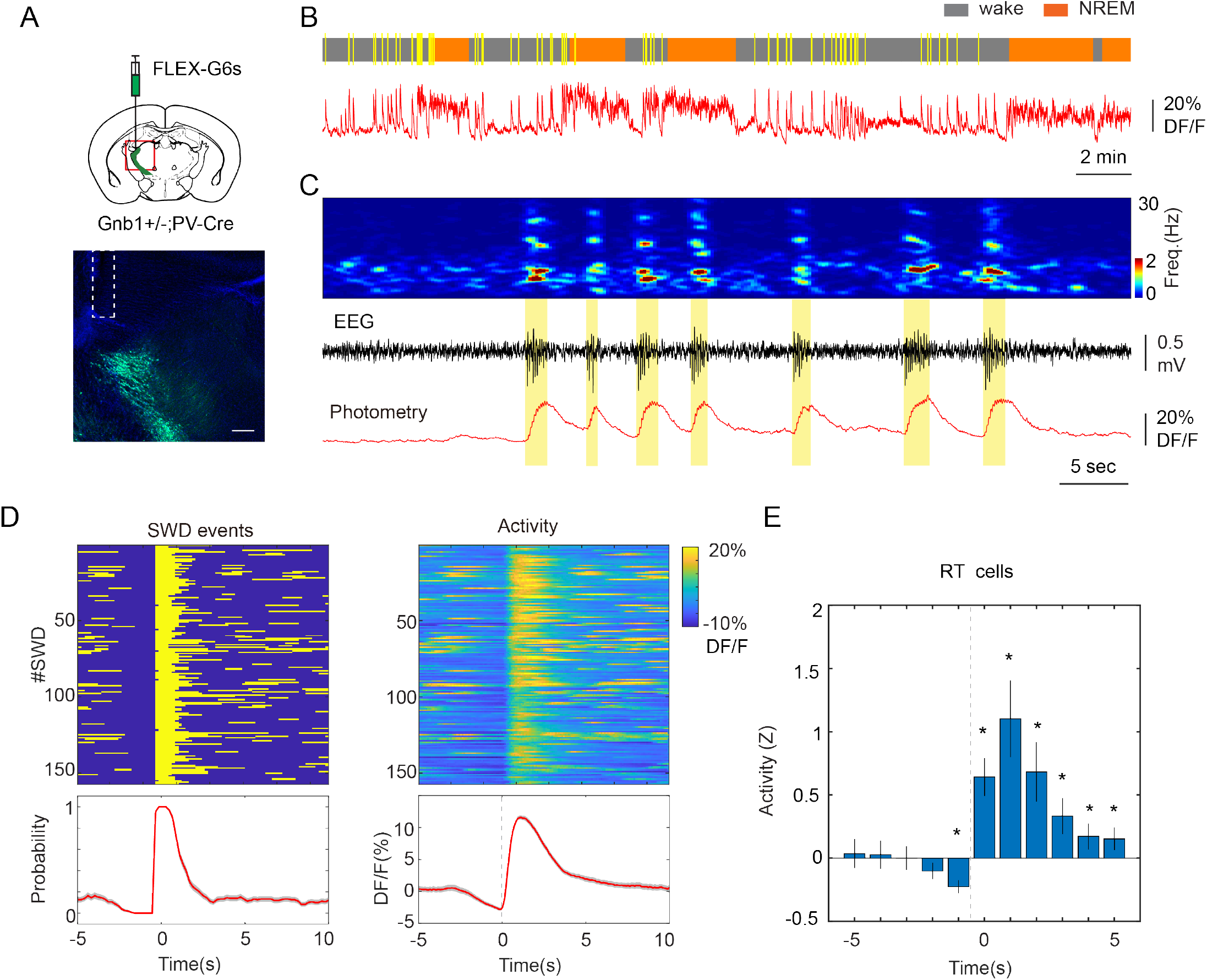
Activity of RT cells correlates with SWD in Gnb1 mice. **A,** Schematic of experimental design. AAV-FLEX-GCaMP6s was unilaterally injected in the RT (Atlas of a brain section, AP - 1.0) of Gnb1^+/-^;PV-Cre mice. Bottom, fluorescence image showing GCaMP6s expression (green) in the RT area (red box above). The white dash line indicates the placement of optic fiber. Blue, DAPI. Scale bar, 200 um. **B,** A representative recording session showing neural activity in RT cells of a Gnb1 mutant mouse during wake and sleep states. **C**, A representative example showing neural activity in RT cells of a Gnb1^+/-^;PV-Cre mouse during SWD. From top to bottom, EEG spectrogram (0-30 Hz), EEG trace, and photometric signal. **D**, Left, an example of aligned SWD events in a recording session (1 hour). Right top, color-coded calcium signals aligned to the onset of SWD events in the recording session shown on Left. Right bottom, averaged calcium signals across trials. Shading, SEM. **E**, Quantitation of calcium activity of RT cells before, during, and after SWD. Datapoints at Time −5 were used as baseline for statistical comparison. The dash line indicates the onset of SWD events. Data represent Z values, normalized to the average DF/F during wake periods (8 recording sessions from 5 Gnb1^+/-^;PV-Cre animals, * P<0.05, paired t-test).

Another notable phenomenon in photometry recording is oscillatory calcium signals during SWD observed both in TC and RT cells (**Figure 6**). This oscillation, indicative of synchronized burst firing, was not observed in the time window prior to SWD. Spectral analysis revealed that the peak frequency of calcium oscillation centered around 7 Hz (**Figure 6B, 6E**), matching the peak frequency of SWD. These results suggest that despite opposite tonic activity, both TC and RT cells display increased burst firing during SWD. Together, our data indicate that the thalamocortical circuits are involved in SWD of *Gnb1* mutant mice.

**Figure 6.**
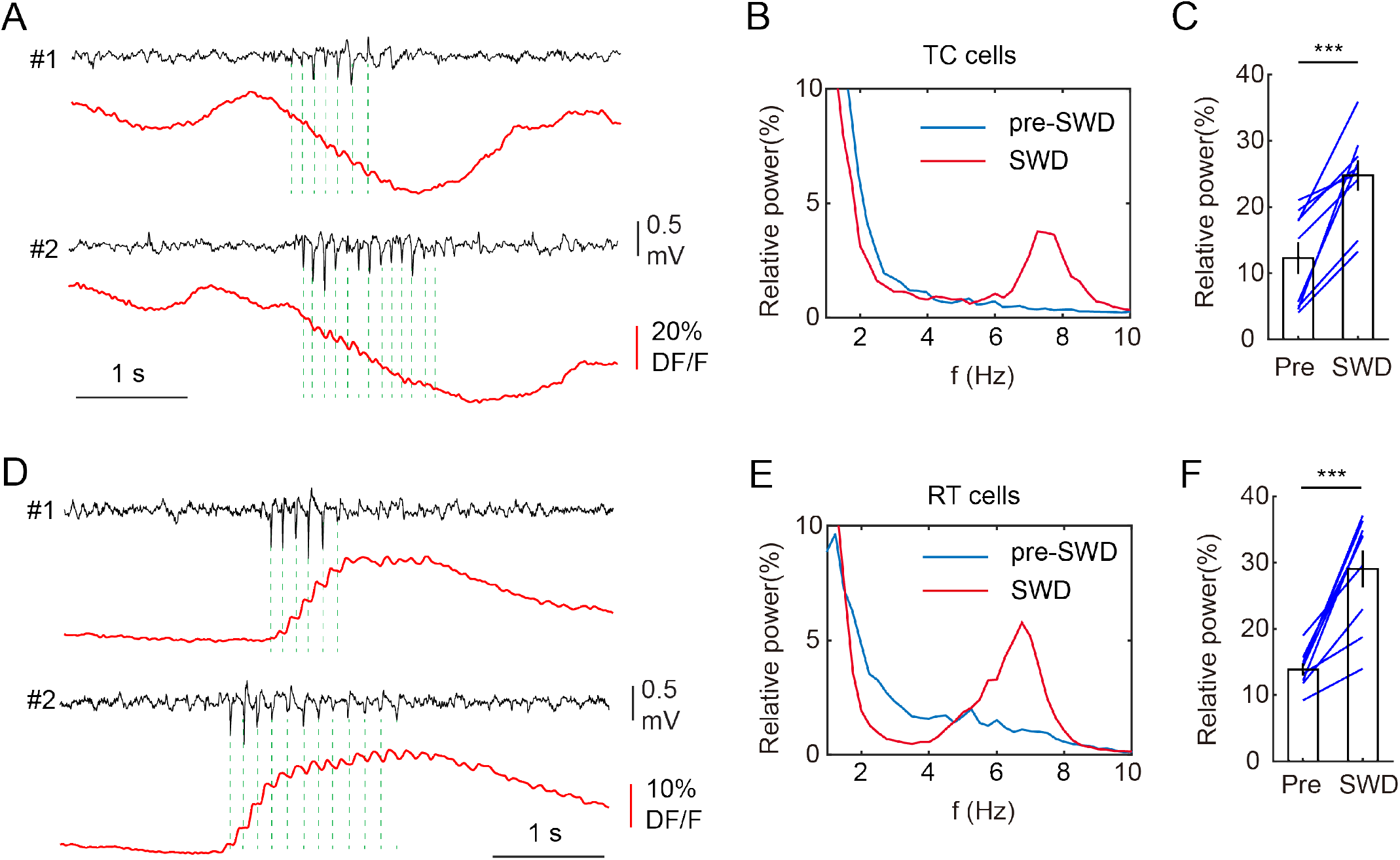
Oscillatory calcium signals in TC and RT cells during SWD. **A,** Two representative SWD events showing SWD-coupled calcium oscillation in TC cells. **B**, Spectral analysis of calcium signals of TC population in the 2-s time window prior to SWD (pre-SWD) and during SWD in a recording session. Power was normalized to the total power in each dataset. **C**, Quantitation of relative power of calcium signals in TC cells during the periods of pre-SWD and SWD (N = 9 recording sessions from 4 Gnb1 mice. *** P<0.001, paired t-test). **D**, Two representative SWD events showing SWD-coupled calcium oscillation in RT cells. **E**, Spectral analysis of calcium signals of RT population in the 2-s time window prior to SWD (pre-SWD) and during SWD in a recording session. Power was normalized to the total power in each dataset. **F**, Quantitation of relative power of calcium signals in RT cells during the periods of pre-SWD and SWD (N = 9 recording sessions from 5 Gnb1^+/-^;PV-Cre mice. *** P<0.001, paired t-test).

### Neural manipulation of thalamic circuits

To further examine the causal effect of thalamic activity on the regulation of SWD, we next performed chemogeneic activation of thalamic cells in Gnb1 mice. To avoid activation of GABAergic RT cells due to viral spread, we crossed Gnb1 mutant mice with VGLUT2-Cre transgenic mice and stereotaxically injected AAV-DIO-hM3Dq-mCherry in the VPM of Gnb1+/-;VLGUT2-Cre (**Figure 7A**). After two weeks of recovery, we treated animals with clozapine-N-oxide (CNO, 2 mg/kg) or saline while recording EEG and EMG. We found that chemogenetic activation of TC cells significantly increase the number of SWD in Gnb1 mice (**Figure 7B-C**). We did not observe a significant difference in SWD duration between CNO and saline treatments (**Figure 7C**). This result indicates that thalamic activity is sufficient to facilitate the SWD in Gnb1 mice.

**Figure 7.**
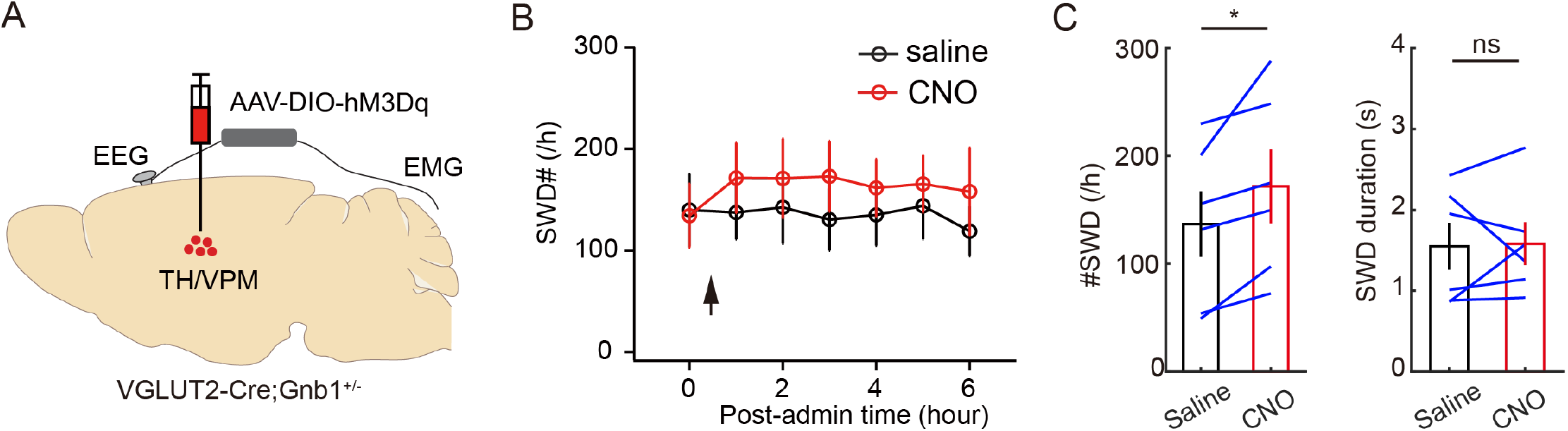
Neural manipulation of thalamocortical circuits in Gnb1 mice. **A,** schematic of experimental design. AAV-DIO-hM3Dq was unilaterally injected in the thalamus of Gnb1^+/-^; VGLUT2-Cre mice. **B**, The number of SWD events per hour in Gnb1 mice following CNO (2 mg/kg, i.p. red circles) and saline (black circles) treatments. Data were normalized to the amount of wake time. Time 0 represents data over 1 hour prior to the treatments (black arrow). **C**, Left, quantitation of SWD number over the first 3 hours following CNO and saline treatments. Right, quantitation of SWD duration over the first 3 hours following CNO and saline treatments. Each line represents one mouse (N = 6 animals, * P<0.05, ns, no significance, paired t-test).

## DISCUSSION

Thalamocortical circuits that underlie sleep spindles under physiological conditions are widely recognized as the main generators of SWD under pathological conditions^2–4,28^. This hypothesis views SWD as a transformation of sleep spindles. In this study, we provide *in vivo* evidence to support that indeed the thalamocortical circuits are involved in SWD in a *Gnb1* mouse model of absence epilepsy. However, our correlation analysis between SWD and sleep demonstrates that SWD predominately occur during wakefulness rather than sleep states in three independent mouse models of absence epilepsy - *Gnb1, Stxbp1* and *Gria4*. A prior EEG study in GAERS rats also shows that the most possible transitions to SWD are from active wakefulness^14^. Spindles, on the other hand, are physiological oscillatory events during NREM sleep^31^, an observation that creates a puzzle when attempting to link sleep spindles to SWD directly. We speculated that some signals happening during wakefulness but not sleep states drive the activity in the thalamocortical circuits which lead to SWD under pathological conditions. Indeed, we show that sensory input itself serves as such a signal as sensory stimulation significantly increased the number and duration of SWD in Gnb1 mutant mice (**Figure 3**). Furthermore, activation of thalamic cells can similarly increase SWD number in Gnb1 mice. Thus, we speculated that lack of SWD during sleep could be due to the absence of sensory inputs. Our finding reveals a possible source of thalamocortical activity that underlies SWD generation. It would be interesting to repeat this observation in other mouse models of absence epilepsy and even in clinical studies in epilepsy patients. Once confirmed, this study might provide translational information to treat absence seizures in humans by advising patients to avoid unnecessary sensory stimulation.

The burst activity of thalamic neuronal populations is considered essential for thalamic rhythmogenesis of SWD^32,33^. Most studies are performed in vitro^8,9^ with compromised network integrity and absence of behavioral relevance. Recently, some *in vivo* electrophysiological studies have investigated thalamic activity during SWD but reported mixed results. For instances, Makinson et al. showed decreased firing in thalamic cells prior to the SWD onset, and slightly increased firing during SWD in *Scn8a* mutant mice^29^. Contradictorily, McCafferty et al. reported decreased total firing in TC cells during SWD in GAERS rats, which was comprised of decreased tonic firing and but increased burst firing^7^. Furthermore, McCafferty et al. also showed increased total and burst firing in RT cells during SWD^7^. Using fiber photometry recording, here we show decreased total ictal activity in TC cells and increased total ictal activity in RT cells (**Figure 4, Figure 5**) during SWD of *Gnb1* mice. Since TC cells receive inhibitory input from the RT cells, the opposite ictal responses between RT and TC cells are expected. Interestingly, our photometry data also revealed SWD-coupled calcium oscillation in both TC and RT cells (**Figure 6**), suggesting that synchronized burst firing in both cell populations are increased during SWD. This result is consistent with burst and tonic results reported by McCafferty et al. The difference of TC ictal activity from that previously reported could be caused by different genetic models of epilepsy, or different brain regions recorded. Interestingly, our data also revealed dynamic activity patterns in RT and TC cells in the time periods that proceed the SWD. Specifically, we demonstrate that RT cells display slightly decreased activity and TC cells show increased activity prior to SWD. Based on our data, we propose the following model of SWD generation: the decreased RT activity might enable more sensory-evoked activity in TC cells during wake periods, which leads to hyperactivity in cortical neurons and subsequently generation of a SWD event in mutant animals. Once a SWD is generated, it activates RT cells, which strongly inhibits TC cells to terminate the SWD event. More *in vivo* studies in other models of absence epilepsy particularly concurrent recordings of cortical and thalamic cells will provide further insight into the cellular mechanisms underlying SWD.

## ACKNOWLEDGEMENTS

We thank Charles Zuker at Columbia University for his support in the early stage of the study. We thank Michael Boland at Columbia University for providing *Stxbp1* mice. We thank Xinyue Chen, Christopher Makinson, and Michael Boland for helpful discussions. This work was supported by startup funds from Columbia University to Y.P., and by Columbia University Precision Medicine Initiative to Y.P.

## AUTHOR CONTRIBUTIONS

S.T., F.Z., and Y.P. designed the study, carried out the experiments, and analyzed data. B.M. performed EEG recording and data analysis. E.Z. performed histology and data analysis. Y.P. and W.N.F. wrote the paper.

## METHODS

### Animals

All procedures were carried out in accordance with the US National Institute of Health (NIH) guidelines for the care and use of laboratory animals, and approved by the Animal Care and Use Committees of Columbia University. Both male and female adult mice which are older than 10-16 weeks of age were used for all experiments. The following mouse lines were used in the current study: C57BL/6J (JAX 000664), VGlut2-IRES-Cre (JAX 028863), PV-Cre (JAX 017320). Gnb1 mice were generated using CRISPR/Cas9 as described previously^13^. The Gnb1 mice used in EEG recordings are on the C57BL/6NJ background. For fiber photometry and chemogenetic manipulation, Gnb1 mutant mice were crossed with Cre lines and F1 mice were used. Stxbp1 mice (JAX 006381) were maintained on a C57BL/6 background^22^. Gria4 mice were maintained as a homozygous colony on a C57BL/6J background^26^. Mice were housed in 12-hour light-dark cycles (lights on at 07:00 am and off at 07:00 pm). For Gnb1 genotyping, DNA was extracted from tail or ear tissue, and PCR was performed, using the KAPA Mouse Genotyping Standard Kit (KAPA Biosystems). The following primers were used for PCR. Fwd: CGAGCATTGAGATCCTCTTTCT; Rev: GTCATCATTGCTCCATCAACAG. The restriction enzyme HinfI was used to distinguish WT from Gnb1K78R/+ mice.

### Viral constructs

AAV1-Syn-FLEX-GCaMP6s, AAV9-CamKIIa-GCaMP6s, AAV8-hSyn-DIO-hM3D(Gq)-mCherry were obtained from Addgene.

### Surgical procedures

Mice were anaesthetized with a mixture of ketamine and Xylazine (100 mg kg-1 and 10 mg kg-1, intraperitoneally), then placed on a stereotaxic frame with a closed-loop heating system to maintain body temperature. After asepsis, the skin was incised to expose the skull and a small craniotomy (~0.5 mm in diameter) was made on the skull above the regions of interest. A solution containing 50-200 nl viral construct was loaded into a pulled glass capillary and injected into the target region using a Nanoinjector (WPI). For EEG and EMG recordings, a reference screw was inserted into the skull on top of the cerebellum. EEG recordings were made from two screws on top of the cortex 1 mm from midline, 1.5 mm anterior to the bregma and 1.5 mm posterior to the bregma, respectively. Two EMG electrodes were bilaterally inserted into the neck musculature. EEG screws and EMG electrodes were connected to a PCB board which was soldered with a 5-position pin connector. All the implants were secured onto the skull with dental cement (Lang Dental Manufacturing). After surgery, the animals were returned to home-cage to recover for at least two weeks before any experiment.

For chemogenetic activation, 200 nl AAV8-hSyn-DIO-hM3D(Gq)-mCherry was unilaterally injected into the VPM (AP −1.8mm, ML 1.5mm, DV 3.2mm) of Gnb1 mice.

For fiber photometry, 150-200 nl AAV1-FLEX-GCaMP6s or AAV9-CaMKII-GCaMP6s was unilaterally injected in the RT (AP −0.9mm, ML 2.0mm, DV 3.5mm) or VPM respectively. An optical fiber (0.2 mm diameter, 0.39 NA, Thorlabs) was implanted 0.2mm above the injection site. The DV Coordinates listed above are relative to the pial surface.

### EEG recording

Mouse seizure and sleep behavior were monitored using EEG and EMG recording along with an infrared video camera at 30 frames per second. Recordings were performed for 24-48 hours (light on at 7:00 am and off at 7:00 pm) in a behavioral chamber inside a sound attenuating cubicle (Med Associated Inc.). Animals were habituated in the chamber for at least 4 hours before recording. EEG and EMG signals were recorded, bandpass filtered at 0.5-500 Hz, and digitized at 1017 Hz with 32-channel amplifiers (TDT, PZ5 and RZ5D or Neuralynx Digital Lynx 4S). For sleep analysis, spectral analysis was carried out using fast Fourier transform (FFT) over a 5 s sliding window, sequentially shifted by 2 s increments (bins). Brain states were semi-automatically classified into wake, NREM sleep, and REM sleep states using a custom-written MATLAB program (wake: desynchronized EEG and high EMG activity; NREM: synchronized EEG with high-amplitude, delta frequency (0.5–4 Hz) activity and low EMG activity; REM: high power at theta frequencies (6–9 Hz) and low EMG activity). Semi-automated classification was validated manually by trained experimenters. For SWD analysis, FFT of EEG was performed using a 1-s sliding window, sequentially shifted by 0.25-s increments. Then, the “seizure”-power (19-23 Hz) was calculated to extract SWD events based on a threshold of 2-3 standard deviations. We chose the 19-23 Hz band to detect SWD based on its clear separation from normal brain oscillatory activities, although the primary spectral band of SWD in mice is around 7 Hz (overlapped with theta oscillations during REM sleep or active periods). Two SWD events were merged into one event if their interval was shorter than 1 sec. Any SWD event with duration less than 0.5 sec was removed for analysis. Algorithm-detected SWD events were further reviewed by trained experimenters. To correlate SWD events with sleep/wake states, each SWD event was aligned to its onset and the probability of brain states 30s before and 30 s after event onset was calculated.

To analyze the effect of SWD on locomotion, a custom-built MATLAB software was developed to perform real-time video-tracking while simultaneously conducting EEG recording. The software synced video-taping with EEG recording through an API provided by TDT. An infrared camera was used to track the body position (the center of the whole body) of a mouse by subtracting each video frame from the background image, captured in the absence of the mouse. The animal’s movement was calculated as the pixel distance between body positions dividing by the time. Then, movement during SWD periods was averaged for each SWD duration or each inter-SWD interval, and further normalized for each animal to the average movement over the whole recording session.

### Fiber Photometry

Fiber photometry recordings were performed essentially as previously described^34^. In brief, Ca2+ dependent GCaMP fluorescence were excited by sinusoidal modulated LED light (465 nm, 220 Hz; 405nm, 350 Hz, Doric lenses) and detected by a femtowatt silicon photoreceiver (New Port, 2151). Photometric signals and EEG/EMG signals were simultaneously acquired by a real-time processor (RZ5D, TDT, sampling rate of 1017 Hz) and synchronized with behavioral video recording. A motorized commutator (ACO32, TDT) was used to route electrical wires and optical fiber. The collected data were analyzed by custom MATLAB scripts. They were first extracted and subjected to a low-pass filter at 20 Hz. A least-squares linear fit was then applied to produce a fitted 405 nm signal. The DF/F was calculated as: (F-F0)/F0, where F0 was the fitted 405 nm signals. Data were smoothed using a moving average method over 0.1s. To compare activity across animals, photometric data were further normalized using Z-score calculation in each mouse.

### Chemogenetic manipulation

After habituation for 12 h in the recording chamber, mice expressing hM3Dq were injected with saline (day 1) and CNO (day 2, 2 mg/kg body weight) intraperitoneally (i.p.) at the same time of the day. EEG recording started at least 1 h before saline injection and lasted 24 h after CNO injection. Data in the time window (0 – 6 h after CNO or saline injection) were used for SWD analysis.

### Histology

Viral expression and placement of optical implants were verified at the termination of the experiments using DAPI counterstaining of 100 μm coronal sections (Prolong Gold Antifade Mountant with DAPI, Invitrogen). Images were acquired using a Zeiss 810 confocal microscope. Cell numbers were counted manually in ImageJ.

### Statistics

No statistical methods were used to predetermine sample size, and investigators were not blinded to group allocation. No method of randomization was used to determine how animals were allocated to experimental groups. Mice in which post hoc histological examination showed viral targeting or fiber implantation was in the wrong location were excluded from analysis. Paired and unpaired T-test were used and indicated in the respective figure legends. All analyses were performed in MATLAB. Data are presented as mean ± s.e.m.

## Code availability

Custom scripts for EEG/EMG and behavioral analysis are available from the corresponding author upon reasonable request.

## Data availability

All data supporting the findings of this study are available from the corresponding author upon reasonable request.

