## Supplemental figures for "Sensory regulation of absence seizures in a mouse model of *Gnb1* encephalopathy"

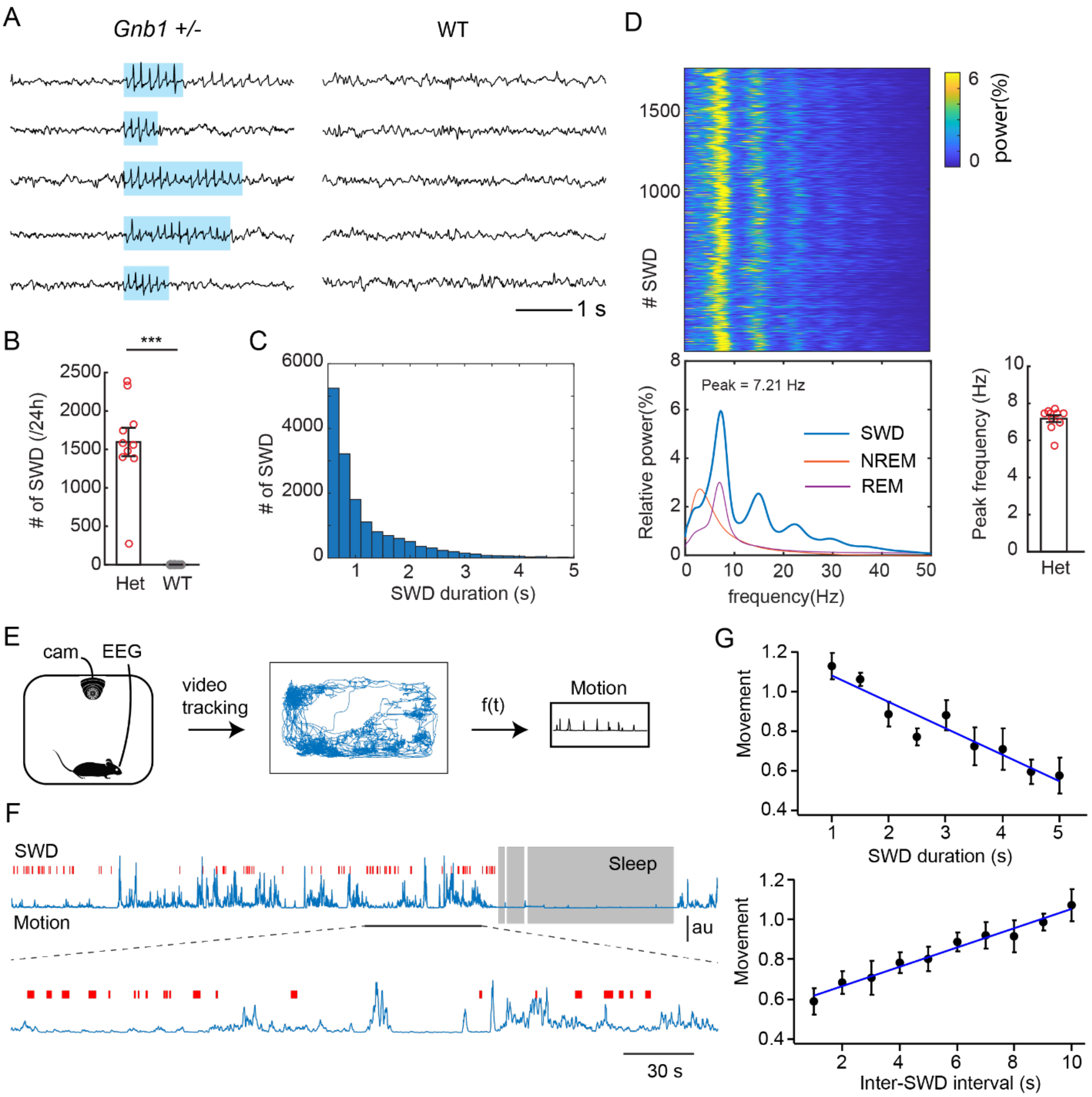


**Supplementary Fig. 1 Characterization of SWD in Gnb1 mice. A**, Representative EEG traces in Gnb1 mutant and wildtype mice. Shading indicates the detected SWD events. **B**, Quantitation of SWD number in Gnb1 heterozygous (Het) mutant and wildtype (WT) mice over 24 hours. Each datapoint represents one mouse (N = 10 mutant, N = 9 wildtype; *** P<0.001, unpaired T-test). **C**, Distribution of SWD durations. Events shorter than 0.5s were excluded. Data from 10 Gnb1 mutant mice. **D**, Left top, a representative example of EEG spectral power (0 – 50 Hz) during SWD in a Gnb1 mutant mouse. Each row represents one SWD event (total 1745 events captured over 24 hours). Left bottom, averaged relative power across SWD events shown above.

The power was normalized to the total EEG power. Relative power of NREM and REM sleep in the same recording session were also plotted as references. Right bottom, quantitation of peak frequency of SWD. Each datapoint represents one mouse (N = 10 Gnb1 mutant mice). **E,** Schematic of experimental design for video tracking. Mice were EEG-video recorded in a behavioral chamber, their locations were tracked by video-processing, and locomotion was calculated from time-serial locations. Middle, A representative example of video-tracking over 30 minutes in a Gnb1 mouse. **F**, A representative example of animal's locomotion (blue trace) correlated with SWD events (red lines). The area with the black underline is enlarged in the bottom. Sleep period (gray) was classified from EEG and EMG data. **G**, Top, correlation analysis between SWD durations and relative locomotion during SWD. Bottom, correlation analysis between inter-SWD intervals and relative locomotion. The blue line indicates linear fitting. Locomotion was normalized to the average of moving velocity during wakefulness in each mouse (N = 8 Gnb1 mutant mice).

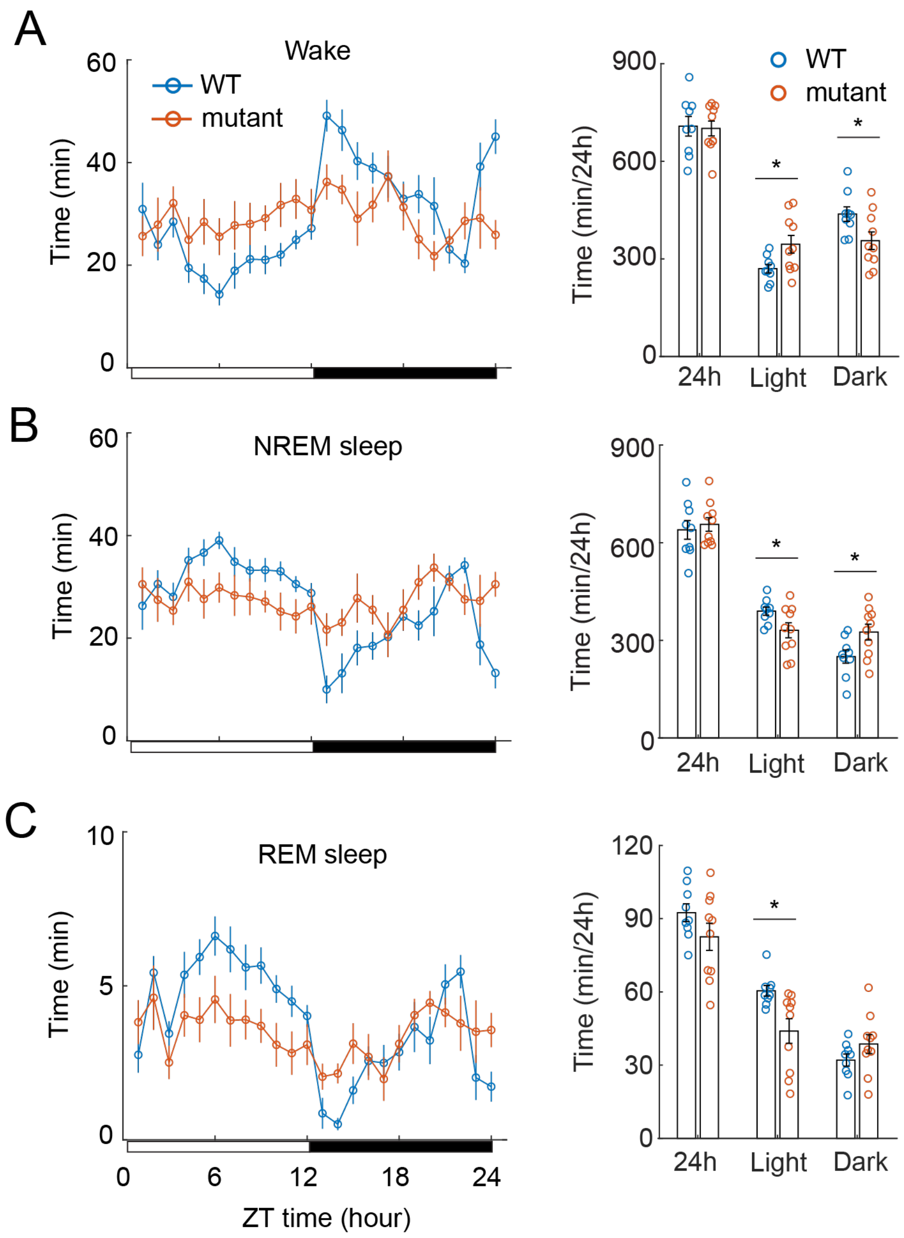


**Supplementary Fig. 2 Sleep patterns in Gnb1 mice. A**, Left, time of wake per hour in Gnb1 mutant and wildtype mice. Right, quantitation of total durations of wake in 24 hours, 12-h light, and 12-h dark cycles in Gnb1 and wildtype mice. **B**, Left, time of NREM sleep per hour. Right, quantitation of NREM sleep time. **C**, Left, time of REM sleep per hour. Right, quantitation of REM sleep time (N = 10 for Gnb1 mutants, N = 10 for wildtype. * P<0.05, unpaired t-test).

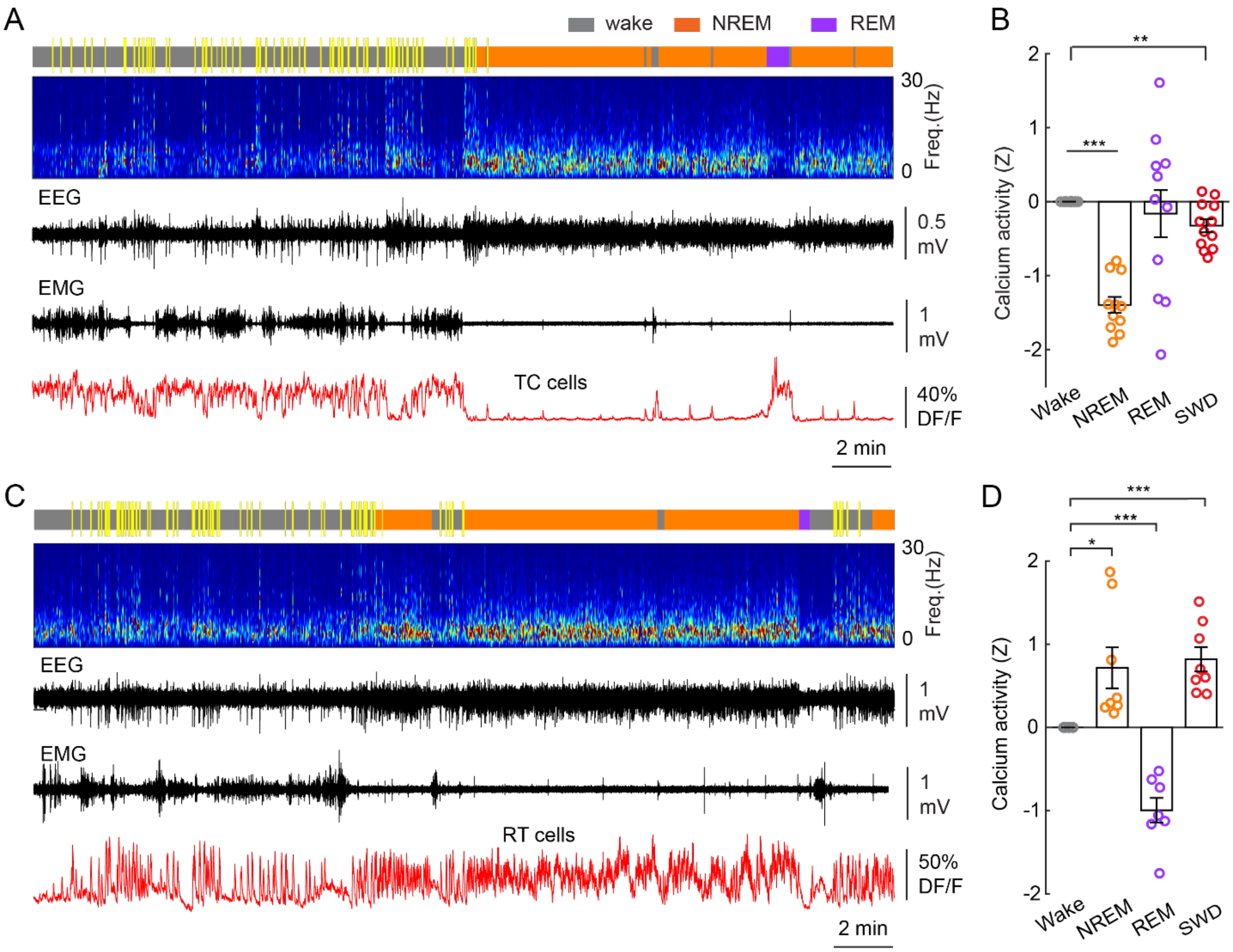


**Supplementary Fig. 3 Activity of thalamic cells during sleep. A,** A representative recording session showing neural activity in TC cells of a Gnb1 mutant mouse during wake and sleep states. From top to bottom, brain states (gray, wake; orange, NREM sleep; purple, REM sleep; yellow, SWD), EEG spectrogram (0-30 Hz), EEG trace, EMG trace, and photometric signal. **B**, Quantitation of calcium activity of TC cells during different brain states (12 recording sessions from 4 animals, ** P<0.01, *** P<0.001, paired t-test). **C**, A representative example showing neural activity in RT cells of a Gnb1$^{+/-}$;PV-Cre mouse during wake and sleep states. **D**, Quantitation of calcium activity of RT cells during different brain states (8 recording sessions from 5 animals, * P<0.05, ** P<0.01, *** P<0.001, paired t-test). Data represent Z values, normalized to the average DF/F during wake periods.
